# SIMple: A fibre-based platform for accessible structured illumination microscopy

**DOI:** 10.1101/2025.09.11.675556

**Authors:** Rebecca M. Mcclelland, Edward N. Ward, Francesca W. van Tartwijk, Stephen Devlin, Junqing Wang, Clemens F. Kaminski

## Abstract

Structured illumination microscopy can be used to achieve optical sectioning and super resolution in fluorescence images, reducing out-of-focus light and increasing the resolution beyond the diffraction limit, without the need for specialised detection optics. However, the complex illumination path is difficult to build and align. We present an illumination path based on fibre-optic components for both splitting and phase-shifting the illumination light. This enables a *SIMple* and compact “Plug&Play” modality which substantially reduces the time and alignment required when adding the optics to an existing widefield instrument. The system is capable of optical sectioning imaging at camera-limited frame-rates using multiple excitation wavelengths simultaneously, as demonstrated by imaging fixed and live biological samples at 561 and 491 nm. Super-resolution imaging of fixed samples on a very compact, self-contained microscope is also demonstrated: illumination is coupled in by fibre to a lightweight frame with dimensions of just 300 × 450 × 300 mm^3^, enabling easy transportation and use in laboratories with limited space. Characterisation of the system using bead analysis shows a resolution of 168 and 172 nm at 491 and 561 nm, respectively, an improvement by a factor of 1.91 and 1.92 compared to widefield, with a field of view of 100 × 100 µm^2^.

## 1. Introduction

Structured illumination microscopy (SIM) is a fluorescence microscopy technique for super-resolution (SR) and optical sectioning (OS) imaging, whereby multiple images of the sample are captured whilst illuminated with different patterns of excitation light. These images can then be post-processed into high-resolution and/or high-contrast images. OS-SIM provides an alternative to laser scanning confocal microscopy, offering similar sectioning capabilities but with reduced photodamage and faster imaging speeds [1]; SR-SIM offers up to a twofold increase in resolution over diffraction-limited imaging. Unlike other SR techniques, such as stimulated emission depletion microscopy and stochastic optical reconstruction microscopy, SR-SIM provides fast image acquisition at low illumination powers that minimise photobleaching and photodamage. This makes it ideal for observing processes such as organelle dynamics in live cells over long durations.

Since the first implementations of SIM [2–6], extensive development has improved imaging performance in terms of resolution, acquisition speed and imaging depth. These advancements have been driven by design and hardware improvements in the optical path as well as new illumination pattern schemes [7–13] and image reconstruction methods [14–19].

Conventional SR-SIM achieves an increase in resolution in two dimensions by using 9 frames of sinusoidal, striped excitation patterns, comprising three rotations each shifted laterally in 3 steps (Fig. 1). The spatial frequency of the excitation pattern mixes with the spatial frequencies of the sample, resulting in the formation of Moiré fringes. These contain high-frequency information about the sample which could not be captured by the diffraction-limited microscope. This high-frequency information can be extracted using the 9 raw images to reconstruct a single SR image. This can, theoretically, achieve a resolution improvement of 2× that of a standard widefield instrument. However, in practice, a resolution increase of 1.7-1.9× is often used to compensate for the attenuation of higher frequencies by the optical transfer function (OTF) and the achievable modulation depth of the illumination patterns.

**Fig. 1.**
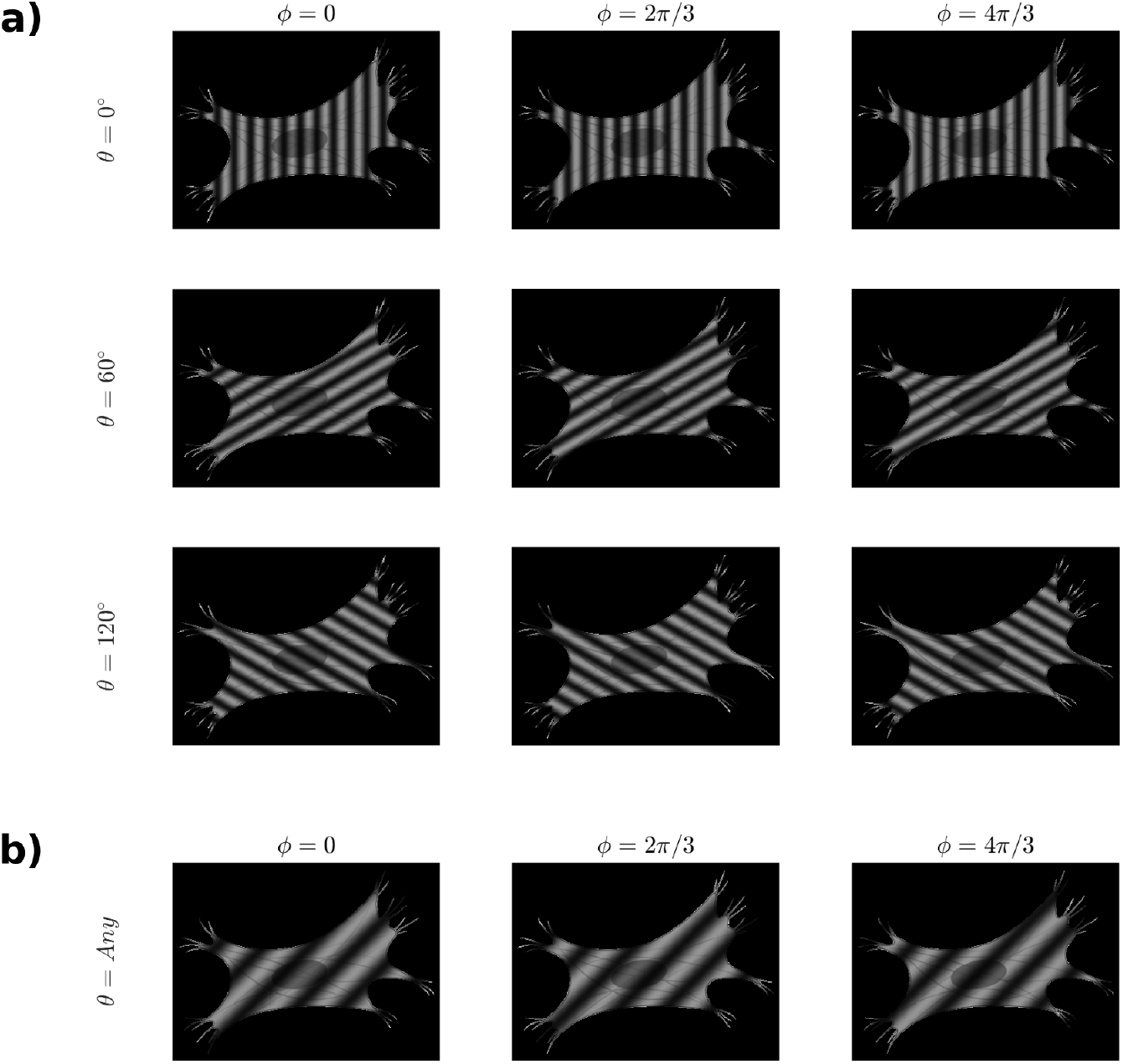
Raw images required for a) SR-SIM and b) OS-SIM. a) For SR-SIM, illumination patterns must be phase-shifted (*ϕ*) 3 times and rotated (*θ*) 3 times. b) For OS-SIM, only three phase-shifted illumination patterns are required, and they can be at any orientation. The spatial frequency of the stripes is lower than for SR-SIM.

Alternatively, OS-SIM can reduce out-of-focus signal in an image using only 3 frames, where the striped illumination pattern is laterally shifted while the orientation remains constant. For optimal sectioning performance, the spatial frequency of this pattern must be approximately half that of the patterns used for SR-SIM (Fig. 1). This is because the aim of OS is to fill the missing cone of frequencies along the *k*_*z*_ axis, rather than to extend the OTF to cover higher frequencies in *k*_*x,y*_ (Appendix A). In this imaging mode, the contrast of the stripe pattern serves as an indicator for whether an area of the sample is in focus, as in-focus areas will be modulated by stripes with a high contrast. The sectioning ability achieved by OS-SIM is similar to that of scanning confocal microscopy [1; 2], but achieves higher image acquisition speeds with lower illumination powers, making it an attractive alternative for live and long-term imaging.

For striped-illumination SR-SIM, four key processes are performed in the illumination optical path to create the excitation patterns: A) an incoming beam is split into two or three beams of equal power for either 2D or 3D SIM, B) the phase difference between the beams is changed to laterally shift the pattern in the sample plane, C) the angle at which the beams cross in the sample is rotated around the optical axis in two steps of *θ* = 60° to rotate the pattern, and D) the beams are relayed to the back focal plane (BFP) of the imaging objective lens, which collimates and overlaps them in the sample plane. The spatial frequency of the pattern is governed by the angle at which the beams leave the objective, which is in turn determined by the location of the beams at the BFP. OS-SIM follows a similar process, although omits step C as only a single pattern orientation is required.

Spatial light modulators (SLMs) and digital micromirror devices (DMDs) can perform steps A, B and C in a single device, with the illumination being relayed to the objective by means of static bulk optics [14; 20–22]. These implementations have the advantage of fast acquisition speeds with sequential imaging of multiple colour channels. However, these devices introduce a number of limitations. The field of view (FOV) is limited by the size of the SLM/DMD active area, and larger devices with the requisite optical flatness are limited in availability. Additionally, the need for sequential multicolour imaging introduces a temporal offset between channels, which prevents reliable imaging of highly dynamic structures in living systems. Finally, the pixelated nature of the devices limits the possible illumination patterns to those that can be represented by an integer number of pixels, making imaging with short wavelengths challenging. Static gratings were used in a similar way in several of the first SIM implementations, but need to be mechanically shifted and rotated to achieve B and C, limiting the imaging speed [3; 4]. Other implementations based on interferometers use beamsplitters to perform step A, and a series of mirrors that are translated or rotated to achieve steps B and C ([16; 23–25]). As beamsplitters and mirrors are achromatic over the visible range, they have the advantage of generating SIM patterns in multiple colours simultaneously. However, these interferometric systems have the drawback of requiring substantial technical expertise in optics to assemble, align and maintain them. Furthermore, as with other systems based on free-space optics, their optical paths can be of substantial size, and they require construction on vibration isolating optical tables.

An alternative approach is to use fibre-optic components for some of the steps. The implementation by Ortkrass *et al*. used beamsplitter cubes, galvanometric mirrors and micro-electromechanical systems (MEMS) actuators for splitting, phase-shifting and switching between pattern orientations, but took advantage of 6 optical fibres to relay light between the two sections of the free-space optical path, in order to overcome the geometry limitations of recombining 6 beams travelling along different paths into a hexagonal configuration at the BFP of the objective [26]. Hinsdale *et al*. used 6 fibre patch cables in a similar fashion, but in combination with Pockels cells, polarising beamsplitters and quarter wave plates for splitting the input light and rotating the illumination patterns. Translation of the patterns was achieved with in-line fibre phase shifters (FPSs) [27]. Hu *et al*. used in-line fibre splitters to split the input beam and fibre switches to direct light to different combinations of outputs for pattern rotation [28]. However, without the ability to shift the phase of the illumination pattern, the technique required non-conventional reconstruction methods. Similarly, Pospíšil *et al*. demonstrated single-colour 2D-SIM using fibre components for splitting the input light and switching between 6 outputs. Pattern phase-shifting was achieved using custom home-built mounts with piezoelectric actuators to physically displace the fibre tips [29].

In this work, we present an optical illumination path that makes SIM imaging more accessible as the optics are easier to assemble and the overall system is more compact and lightweight, without compromising imaging performance. The spacing of the illumination pattern can be changed easily to switch between different SIM imaging modes (e.g. OS and SR) or different objective lenses without any re-alignment of the optics. This is achieved by splitting, phase-shifting and out-coupling the illumination using off-the-shelf, in-line fibre components, and reducing the length of the free-space optical path between the fibre outputs and the tube lens to a compact 300 mm. This concept is implemented in two ways. First, we present an “add-on” OS-SIM module that can be straightforwardly assembled from off-the-shelf fibre components and connected to an existing widefield microscope with minimal optical alignment. Secondly, a SR-SIM instrument is described, which comprises a compact 300 × 450 × 300 mm^3^ frame housing the illumination and detection paths, which can be connected to the fibre circuit to form a complete imaging system. OS-SIM imaging is demonstrated at 491 and 561 nm on fixed and live cell samples and SR-SIM imaging is demonstrated on fixed cells, achieving a resolution of 168 nm, a 1.91× increase over widefield imaging. The potential use of this design to address accessibility limitations and enable imaging experiments with better maintenance of a range of conditions is discussed.

## 2. Materials and methods

### 2.1 SIMple optical setup

The layout of the optical illumination path is shown in Fig. 2a. A 491 nm (Cobolt Calypso, HÜBNER Photonics) and a 561 nm (SLIM-561-150, Oxxius) laser line are coaxially combined and coupled into one of the inputs of a 2 × 2 fibre splitter (TW560R5A2, Thorlabs). This input light is split in a 50:50 intensity ratio between the two outputs, one of which is connected to the fibre phase shifter (FPS, FPS-001, Luna Inc.), which is capable of 20 kHz operation. The phase shift is controlled by an applied voltage sent via a DAQ card (PCIe-6353, National Instruments) used in conjunction with a BNC terminal breakout box (BNC-2110, National Instruments). The FPS output and the second output of the splitter are connected to two of the inputs of a fibre array (Meisu Optics, see Appendix B), which outcouples the light into free space with a separation between the beams that is an integer multiple of the channel spacing, 127 µm, depending on which of the 9 inputs are connected. This allows the illumination to be adjusted for different illumination pattern spatial frequencies or different objective lenses without re-alignment (see Appendix B.2). The fibre type is S405-XP, enabling use of the array across the 400 - 680 nm wavelength range. The fibre components are all pigtailed with FC/APC connectors, and therefore are straightforwardly assembled by connecting with fibre mating sleeves (ADAFC4, Thorlabs), removing the need for any alignment. The fibre outputs at the fibre array emit diverging beams that are collimated by a collimating lens (CL, AC254-030-A-ML, Thorlabs) with focal length *f*_*C*_ = 30 mm. An adjustable pinhole (SM1D12C, Thorlabs) and linear polariser (LPVISE050-A, Thorlabs) are placed at the crossover point of the beams to control the excited FOV and the polarisation of the beams. The beams are then focused onto the BFP of the objective (UPLSAPO60XW, Olympus) with a tube lens (ITL200, Thorlabs). As the separation of the two beams at the exit of the fibre array needs to be magnified onto the objective BFP to achieve the required spacing, a short *f*_*C*_ can be used, making the system compact: the fibre array, collimating lens, pinhole and polariser can be mounted in a 300-mm long cage system (Thorlabs) when the tube lens has *f*_*T*_ = 200 mm (Fig. 2b).

**Fig. 2.**
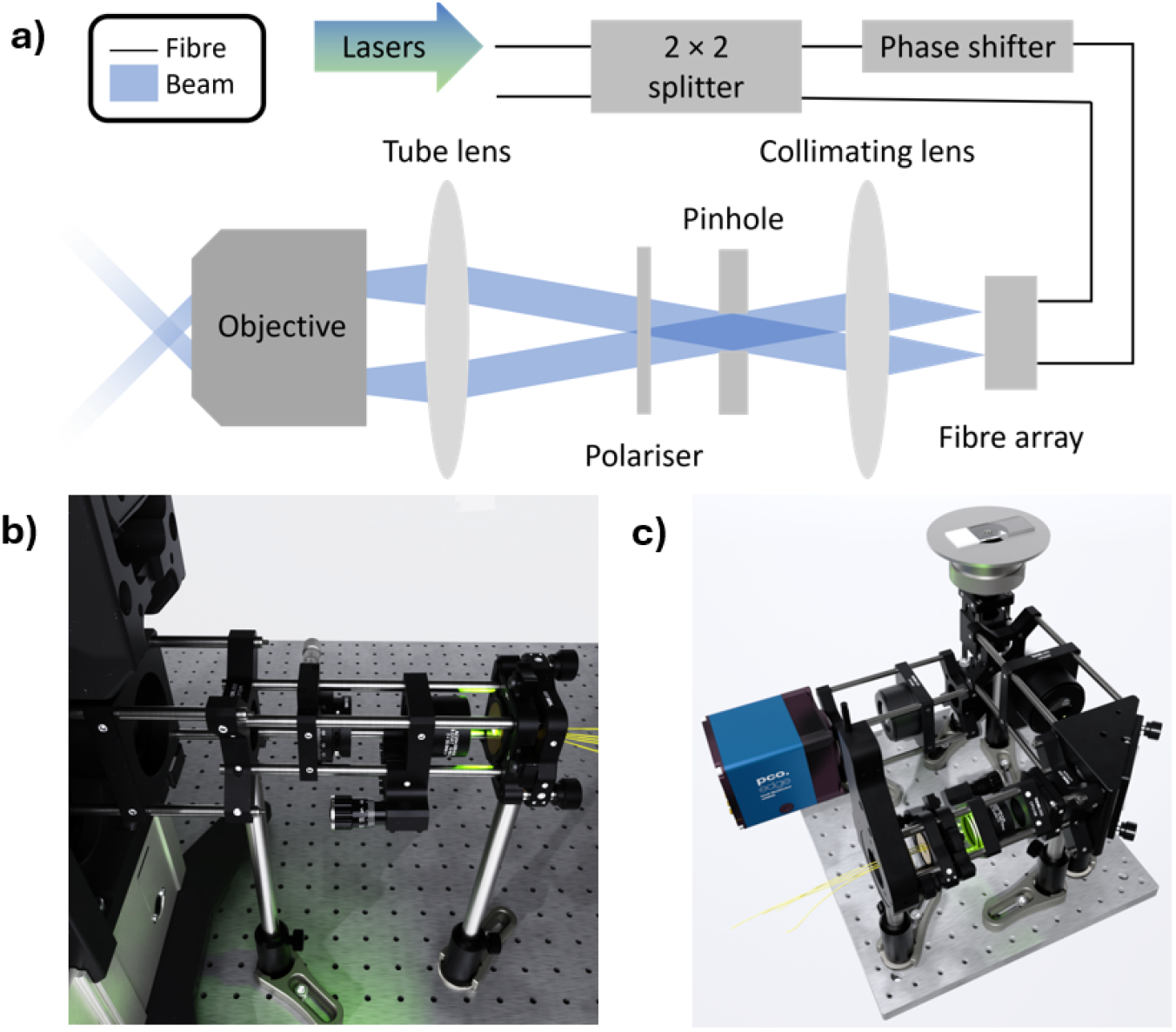
The *SIMple* design’s use of in-line fibre-optic components permits the construction of compact and flexible structured illumination microscopes (SIMs). a) Diagram of the fibre-based illumination path. Multiple input wavelengths are split in a 50:50 intensity ratio, phase-shifted and output to free space by compact, in-line fibre components. A 4f telescope magnifies the illumination beams and relays light to the back focal plane of the objective. b) The fibre-based design applied as an add-on module to an existing widefield microscope to enable OS-SIM. The illumination module has dimensions 60 × 60 × 300 mm^3^ and comprises just 4 optical components: a fibre array mounted in a kinematic mount, a collimating lens, a pinhole and a linear polariser. The use of cage mounted optics reduces the degrees of freedom which greatly simplifies the construction and alignment process. c) Portable stand-alone instrument for OS and SR-SIM imaging, with dimensions of 300 × 450 × 300 mm^3^ (W × L × H). Illumination is coupled in from the fibre circuit via the fibre pigtails of the array. The lightweight system is mounted on a single breadboard for easy transportation by an individual and is compact enough for use in confined spaces.

When attaching the module to an existing widefield microscope via a cage system, minimal alignment is required: the module is positioned in the x-y plane to be aligned with the optical axis of the objective lens, and then the 3 components must be positioned only in the z direction. The use of a dedicated port adaptor would remove the need for any alignment in the x-y plane. On the custom frame, rotation of the patterns for SR-SIM imaging was achieved by placing the fibre array in a rotation mount (PRM1/MZ8, Thorlabs) which was rotated in three 60^°^ intervals, with three phase-shifted images being captured per rotation. Both implementations separated the excitation and detection paths by means of a quad band dichroic mirror (ZT405/488/561/640rpcv, Chroma), and captured images using a 2048 × 2048 pixel sCMOS camera (pco.edge 4.2 bi) with 6.5 µm pixel pitch. Simultaneous multicolour OS-SIM imaging was achieved by means of splitting optics in the detection path (OptoSplit III, Cairn) (Fig. 2b).

### 2.2 Data acquisition and processing

Image acquisition was controlled by custom *Python* software. OS-SIM data were reconstructed by means of machine-learning (ML)-based OS-SIM reconstruction software, previously reported in [17]. SR-SIM data were reconstructed using either the open-source software fairSIM [30], or an adapted MATLAB implementation of an inverse-matrix reconstruction method [31]. Pseudo-widefield comparison images were generated by taking the mean intensity of the 3 or 9 raw frames for OS- and SR-SIM, respectively. The resolution increase of SR-SIM was quantified by measurements of sub-diffraction beads, and image decorrelation analysis (Appendix C) [32].

### 2.3 Sample preparation

Details of sample preparation methods are given in Appendix D.

## 3. Results

### 3.1 Fibre array allows control of the spatial frequency of illumination patterns

To demonstrate the compatibility of the fibre-based illumination module with a range of SIM modalities, the spatial frequency of the sinusoidal stripe pattern produced was measured as a function of the spacing between the fibre array outputs (Appendix B.2). The two outputs of the fibre circuit were connected to two input channels of the fibre array with the appropriate spacing, and images were captured of a monolayer of fluorescent beads with a 60× 1.2NA water immersion objective (UPLSAPO60XW, Olympus) and a 100× 1.4NA oil immersion objective (UPLANSAPO100XO, Olympus) (Fig. 3). The spatial frequencies were calculated from the fast Fourier transforms (FFTs) of the images. The results show that the illumination spatial frequency can be quickly and easily changed by connecting the fibre circuit to different inputs of the fibre array. Additionally, the pattern can be reproduced with a high repeatability as the array can remain stationary while the pattern spacing is changed, and no optical re-alignment or re-programming of any electronic components is required. The range of spatial frequencies that can be generated covers the values required for OS-, SR- and TIRF-SIM (Appendix A) and also the range of values of back aperture diameter for most objectives.

**Fig. 3.**
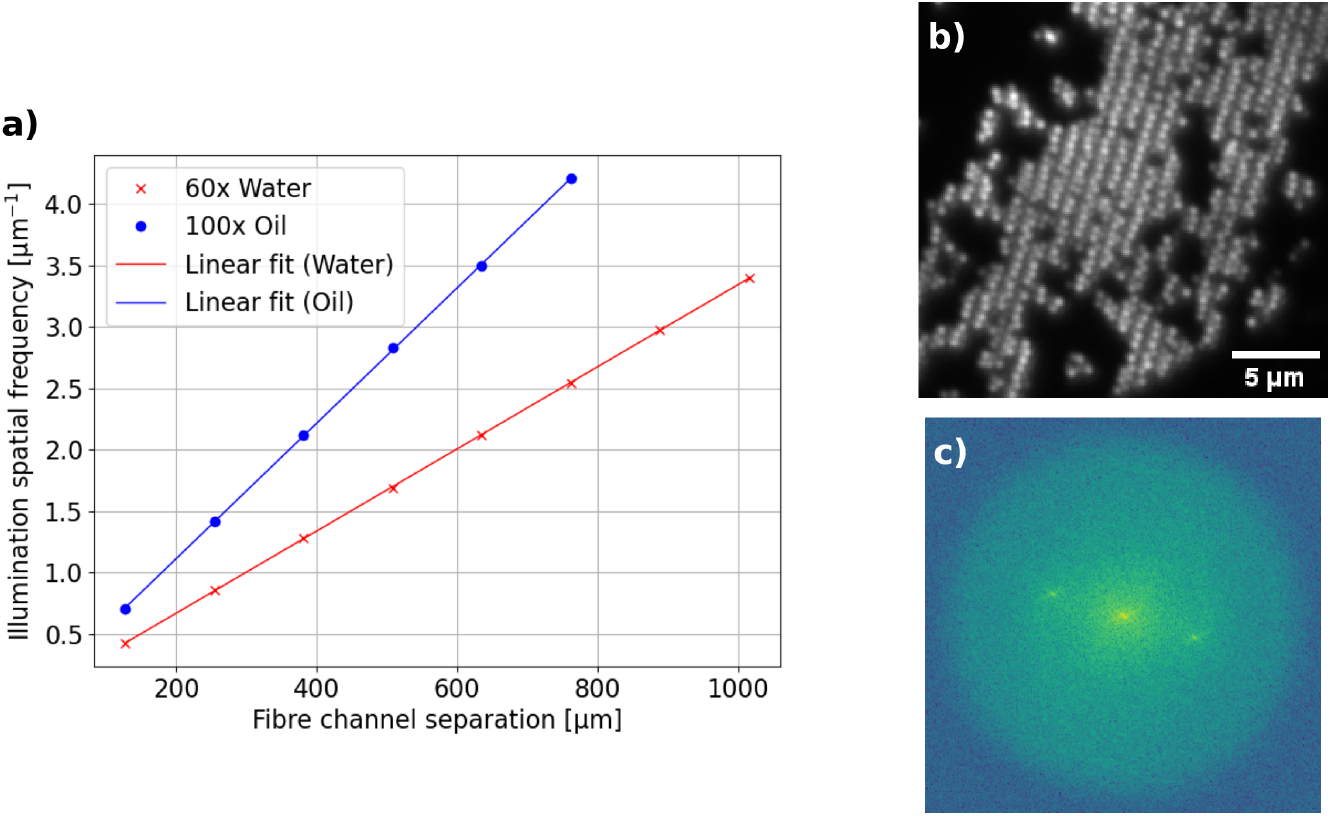
The spatial frequency of the illumination pattern can be controlled by outcoupling light from different channels of the fibre array. a) The spatial frequency of the illumination pattern as a function of separation between the channels used to outcouple light from the fibres, for two different objective lenses (UPLSAPO60XW, Olympus and UPLANSAPO100XO, Olympus). The separation values are integer multiples of the channel spacing (127 µm). Test sample was a monolayer of 250-nm fluorescent beads. The relationship was linear for both objectives. b) Example data of beads imaged with the water objective for a fibre channel separation of 381 µm. Image is a cropped section of the full field of view used. c) The corresponding Fourier transform.

### 3.2 Fibre phase shifter enables OS-SIM pattern generation at 351 Hz

Phase-shifting of the illumination patterns is achieved using the FPS, which is controlled by a DC voltage from the DAQ card. To calibrate the control of the pattern phases, varying voltage values were applied to the FPS and illumination patterns back-reflected from a microscope slide were observed using the camera. The change in phase between frames was calculated and plotted as a function of the change in applied voltage, showing a linear response which was used to calculate the experimental values (Fig. 4a).

**Fig. 4.**
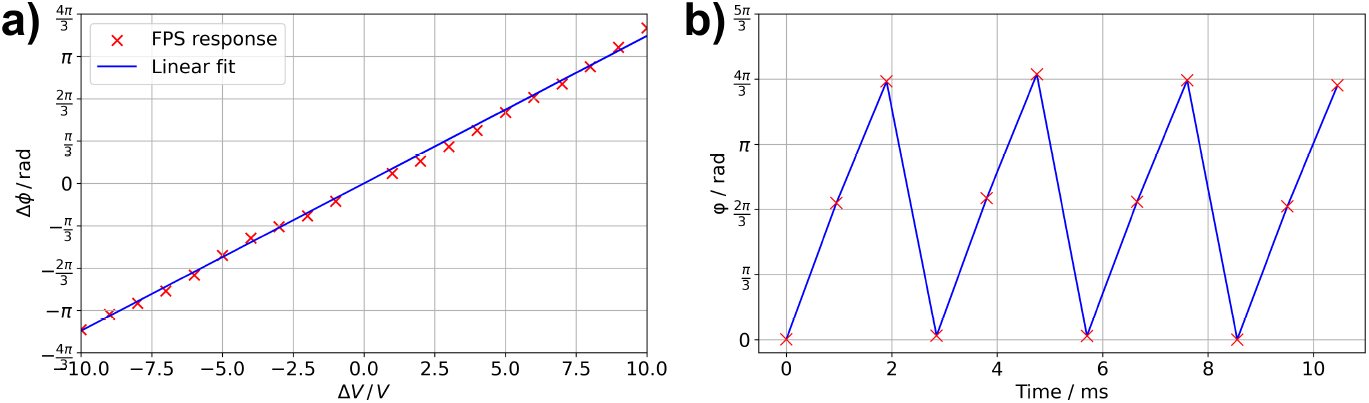
A fibre phase shifter (FPS) can control SIM illumination patterns at camera-limited speeds. a) The phase shift (Δ*ϕ*) generated by the FPS was calibrated as a function of changes in input voltage (Δ*V*), showing a linear relationship. b) Phase-shifts of the SIM patterns can be repeated at >1.053 kHz, here limited by the frame-rate of the camera.

The operation rates of the FPS and DAQ card are 20 kHz and 2.86 MS/s, respectively, suggesting that the image acquisition speed would be camera-limited. To verify this experimentally, a test acquisition was carried out: to maximise the imaging rate, a TTL pulse from the camera was used to trigger the DAQ card, and the FOV was set to 4 × 4 µm^2^ with a 21 µs exposure time. In this configuration, the raw frame rate of the camera was 1053 Hz and calculation of the phase of each image showed correct phase-stepping between each frame (Fig. 4b). A similar acquisition without phase-stepping showed that there was no loss of modulation depth between the two, confirming that the phase-shifts had fully settled between frames. Exact quantification of the actual speed was not considered necessary as at >1053 Hz for raw frames, which corresponds to 351 Hz for OS-SIM reconstructed frames, the limiting factors in a real experiment would be associated with the camera and the sample.

### 3.3 Multicolour optical sectioning imaging enables visualisation of organelle morphology and dynamics

The add-on module was used with an existing widefield microscope to capture images of the cytoskeleton in fixed Vero cells (Fig. 5) and of multiple organelles within live COS-7 cells (Fig. 6), which, compared with widefield data, show an improvement in contrast. Within live cells, this improved contrast allowed the morphology and dynamics of mitochondria and the endoplasmic reticulum (ER) to also be distinguished in the thicker perinuclear region (Fig. 6); this makes the technique ideal for imaging experiments investigating organelle interactions, as it allows contact sites to be observed across the whole cell, not just the flatter periphery, without the trade-off of slow acquisition speeds and/or increased photobleaching when using confocal microscopes [33]. This would be beneficial for imaging experiments where automatic segmentation of organelles is required, enabling clear identification of organelles even with high background signal [34–36].

**Fig. 5.**
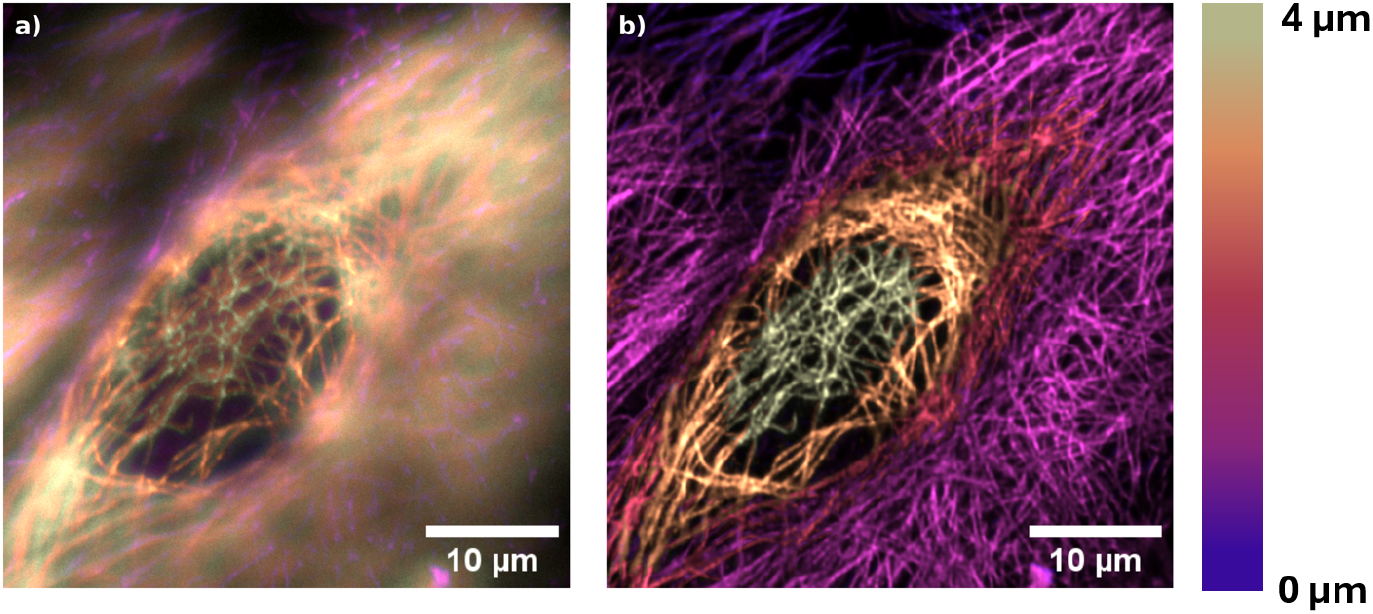
Maximum intensity projections of a) widefield and b) OS-SIM data show a clear reduction of out-of-focus signal in the latter, enabling different heights in the sample to be clearly distinguished. Images are of immunostained *β*-tubulin, illuminated with *λ* = 561 nm. Colourmap represents z position from 0 to 4 µm. Scale bars are 10 µm.

**Fig. 6.**
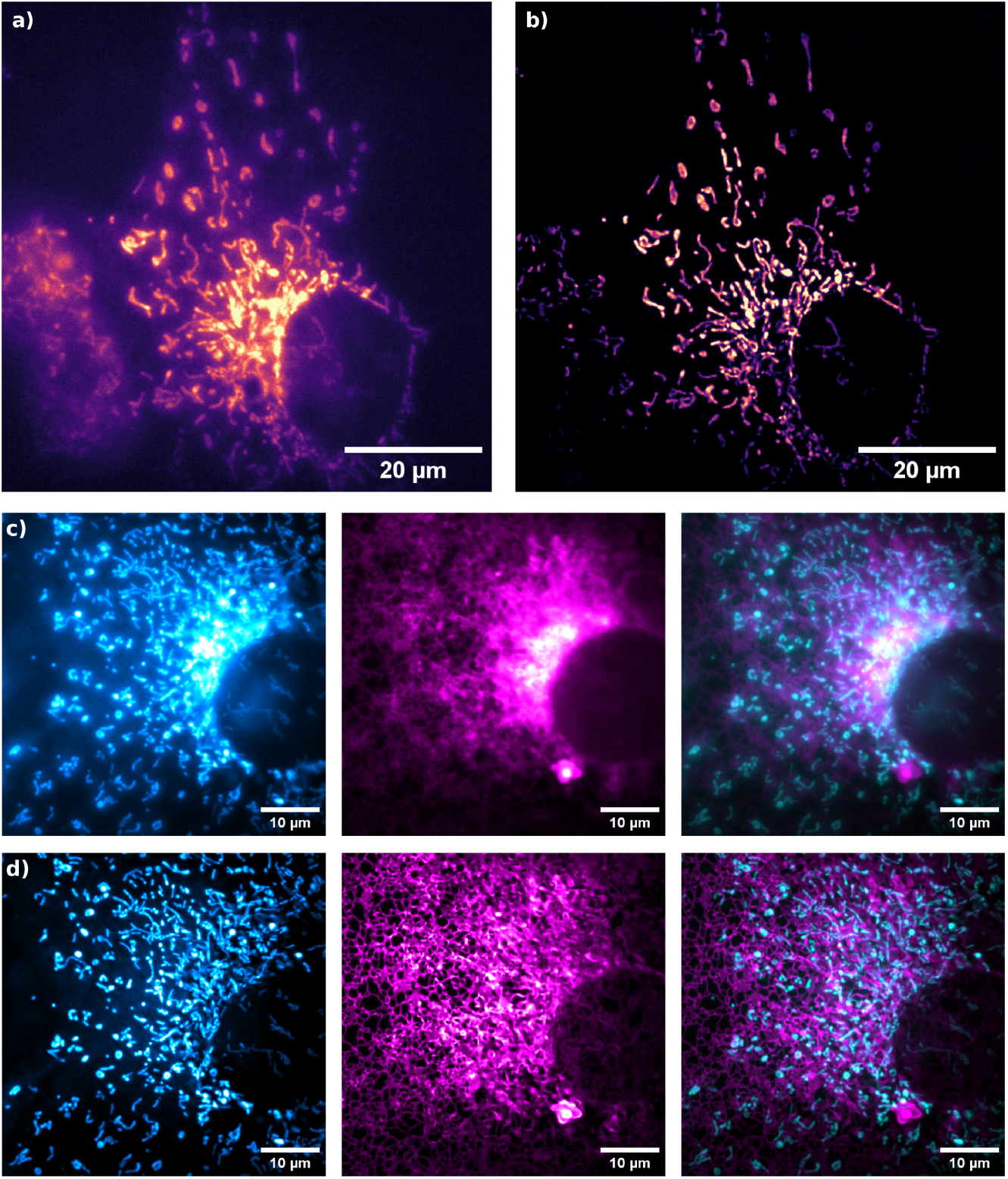
Fibre-based OS-SIM imaging of live COS-7 cells demonstrates reduction in out-of-focus blur when comparing the a) widefield and b) OS-SIM images of mitochondria, allowing individual organelles to be distinguished in the perinuclear region. Scale bars are 20 µm. c) Widefield and d) OS-SIM imaging in multiple channels simultaneously allows clearer visualisation of the morphology and dynamics (see supplementary video) of the mitochondria (cyan) and endoplasmic reticulum (magenta). Scale bars are 10 µm.

### 3.4 Super-resolution imaging shows 1.91 times resolution improvement

A fixed Vero cell sample was imaged with 561 nm illumination to demonstrate SR-SIM with the system (Fig. 7). Two methods were employed to quantify the resolution increase: first, decorrelation analysis of the fixed sample image was performed, revealing a 1.81× resolution increase over widefield imaging, from 363 to 201 nm (Appendix C). Secondly, the full width half maxima (FWHM) of beads were analysed (Fig. 8). For illumination with *λ* = 491 nm, the resolution increased from 321 to 168 nm, a 1.91× improvement over widefield imaging. For *λ* = 561 nm, the resolution increased from 330 to 172 nm, a 1.92× improvement.

**Fig. 7.**
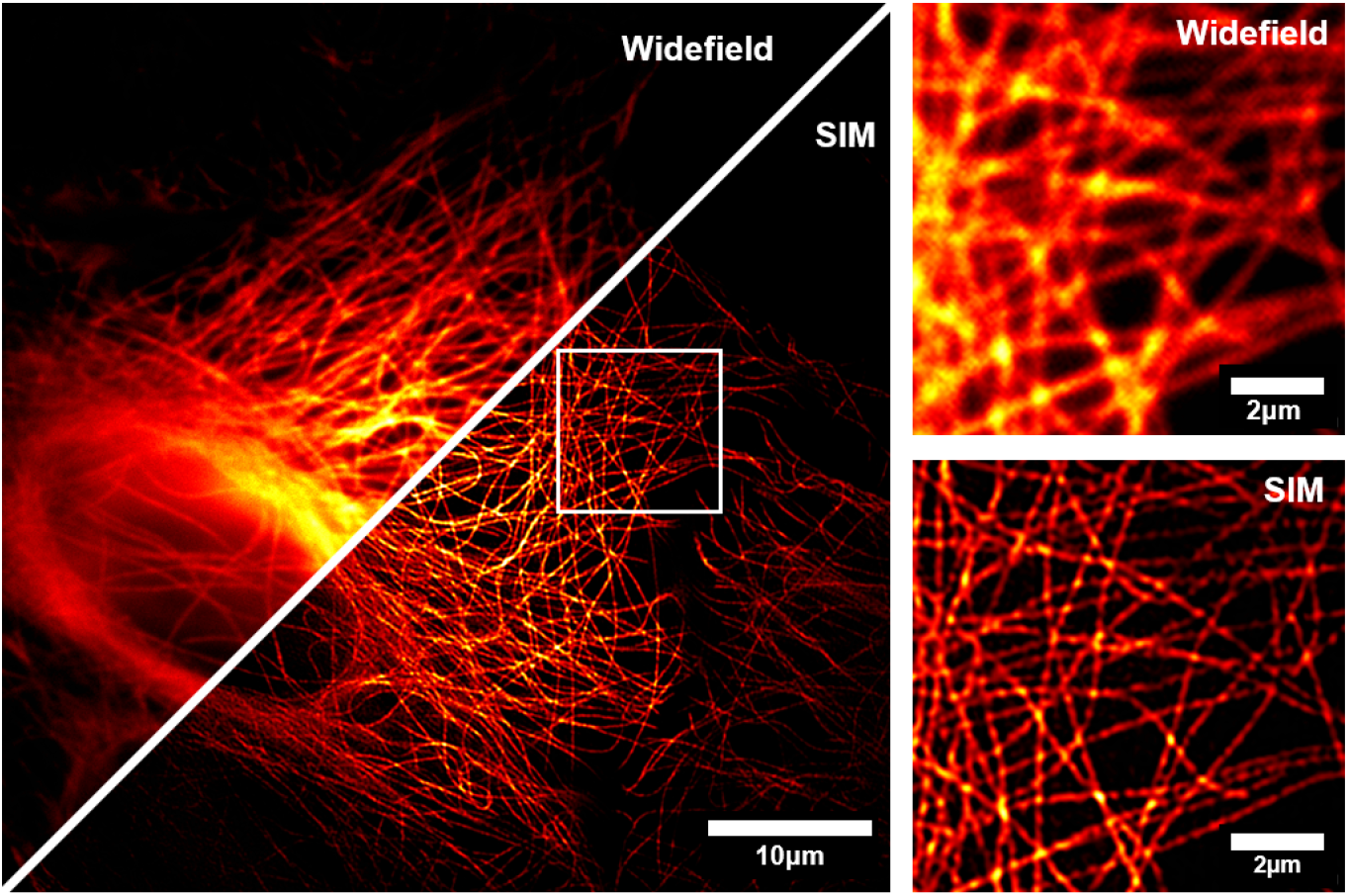
Fibre-based SR-SIM increases the resolution compared to standard widefield imaging. Images are of immunostained *β*-tubulin in fixed Vero cells, illuminated with *λ* = 561 nm. Fourier ring correlation of the image gives a resolution increase of 1.86× compared to widefield imaging. Scale bars are 10 µm (left) and 2 µm (right).

**Fig. 8.**
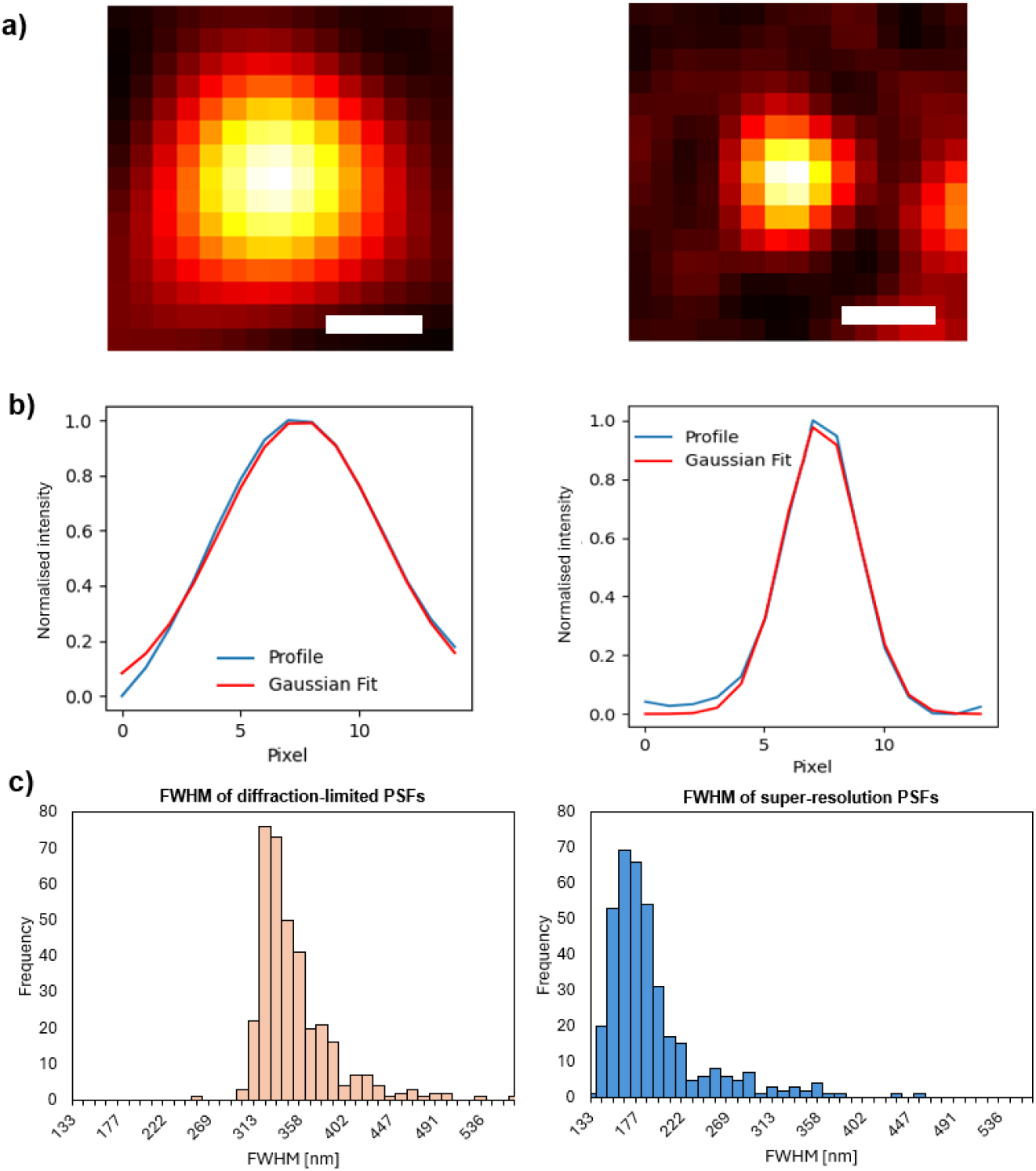
Fibre-based SR-SIM (right) increases the resolution by a factor of 1.92 (*λ* = 561 nm) and 1.91 (*λ* = 491 nm) compared to widefield (left), as quantified by analysis of imaging bead monolayers. a) Representative images captured with *λ* = 561 nm, scale bars are 500 nm. b) Fitted intensity profile of representative bead images. c) Histogram of full width half maxima (FWHM) of the point spread functions (PSFs) of SIM: (*n* = 383) and widefield: (*n* = 359).

## 4. Discussion

Despite the advantages of SR-SIM and OS-SIM, their accessibility to life science researchers remains limited. SIM instruments are typically purchased as complete systems, accessed through imaging centres, or built in-house. Both commercial and home-built instruments are expensive, bulky and require strict environmental control and ongoing maintenance, compromising their use in many laboratories, particularly high biological safety level (BSL) laboratories with strict access limitations. Imaging centres mitigate these issues, but only if samples can be transported safely, and limited instrument availability restricts long-term imaging experiments. Creating a home-built instrument reduces costs and allows customisation, but the considerable time investment and expertise requirement makes this approach infeasible for researchers who do not primarily focus on optical development [37; 38]. More compact systems based on DMDs and MEMS actuators have been proposed [23; 39], but still require specialist optics knowledge to build and a considerable amount of space to house. Low-cost and/or open-source implementations often necessitate a trade-off with imaging performance, and also still require relatively substantial prior knowledge, time commitment and specialist equipment to assemble, which has restricted their adoption by the imaging community [40; 41].

The *SIMple* design addresses these challenges by using easy-to-assemble, off-the-shelf fibre-optic components requiring minimal alignment, minimising the amount of optics expertise required to retrofit widefield microscopes for optical sectioning. This provides an easier and more affordable alternative to scanning confocal or spinning disc add-on modules. Its compact and lightweight design enables a stand-alone instrument that can be carried by a single user, potentially facilitating a SIM version of the *Flamingo* model of portable, reconfigurable and loanable instruments [42; 43]. Currently there are no reports of such systems that can achieve SR imaging. The optical fibre design could allow for easy decontamination of the optics when being used in high BSL laboratories, whilst keeping the delicate laser sources housed far from the sample mounting stage where potential contamination could take place.

SIM is very well-suited for live single-cell imaging, as it combines high resolution and contrast with fast acquisition speeds and low illumination power. However, additional engineering solutions are required to maintain cell health-critical conditions during long-term imaging, such as temperature, humidity and gas concentration [44]. Typically, some apparatus is placed on the microscope stage or around the frame to maintain these conditions [45–50]. However, stage-top devices can be limited by gas leakage or depletion with time, or by thermal contact between the sample and the ambient-temperature objective, necessitating additional custom or commercial equipment to directly control objective temperature [51; 52]. Alternative solutions are compact microscopes placed inside standard, widely-available incubators for effective long-term control [53–56], or commercial solutions where both are integrated in one product. However, these have so far been limited to widefield imaging, which is fundamentally constrained by out-of-focus light and the diffraction limit. Extension to SIM has been hindered by bulky free-space optics and delicate electronics such as the SLM. The *SIMple* microscope’s fibre-based design could address these issues and open up the possibility for easy maintenance of cell health over long-term imaging experiments without the need for custom environmental isolation solutions, such as the subzero temperatures required by cold-adapted Antarctic species [51].

Fibre optics also addresses the drawbacks associated with SLM-based SIM, as they enable simultaneous generation of spacing-optimised patterns in multiple wavelengths, avoiding sequential channel imaging with strongly wavelength-dependent devices. Furthermore, fibres that are suitable for use with the entire visible band are available, meaning that they can work with shorter wavelengths around 400 nm, where the performance of SLMs is limited. The main limitation of the system currently is the SR-SIM acquisition speed. Although camera-limited OS-SIM has been demonstrated, mechanical rotation of the fibre array restricts SR imaging to fixed samples.

This will be addressed in future work by incorporating rapid fibre-based switching between pattern orientations.

## 5. Conclusion

This work presents a new method for generating multicolour SIM patterns simultaneously, based on fibre-optics, which substantially minimises the number of optical components required, making the system very compact. Use of a fibre array with multiple channels enables easy and highly reproducible control of the spatial frequency of the illumination patterns for use with different objectives or different types of SIM imaging, without needing any optical re-alignment. Furthermore, the *SIMple* design minimises the time and optics expertise necessary to build the illumination optical path, as fibre components can be straightforwardly connected to each other. Multicolour OS-SIM imaging of live cells and SR-SIM imaging of fixed cells over a large FOV are demonstrated, with characterisation of the system showing a 1.91× resolution increase compared to standard widefield imaging. The system is compact, robust, transportable and capable of housing the pattern-generating optics separately from the microscope frame by using optical fibres, making it more accessible and opening up the possibility of new types of imaging experiments in unconventional environments.

## Acknowledgments

The authors would like to thank Jacob Lamb for useful discussions about the optical setup. C.F.K. acknowledges funding from the UK Engineering and Physical Sciences Research Council (EPSRC) [EP/L015889/1 and EP/H018301/1], the Wellcome Trust [3-3249/Z/16/Z and 089703/Z/09/Z], the Medical Research Council (MRC) [MR/K015850/1], the UKRI Cross Research Council Responsive Mode (CRCRM) award [MR/Z505341/1], and Infinitus China Ltd. R.M.M. acknowledges funding from the UK EPSRC (EP/S022139/1) and the Alzheimer’s Research UK East Network Centre (ARUK-NC2024-EAST). S. D. acknowledges funding from the EPSRC Centre for Doctoral Training in Sensor Technologies for a Healthy and Sustainable Future [EP/S023046/1]. The authors acknowledge BioRender for its use in the figures.

## Data Availability Statement

Data underlying the results presented, code used for image reconstruction and hardware control and design of custom parts will be made available upon publication and can be obtained from the authors upon reasonable request.

## Appendix A SIM in frequency space

Different spatial frequencies of the illumination patterns are required according to the desired SIM imaging mode, as illustrated in Fig. 9. In the case of widefield imaging, the size of the OTF is limited by the diffraction limit. Notably, the OTF in *k*_*x,z*_ has a toroidal shape with a “missing cone” along the the *k*_*z*_ axis. The loss of this frequency information in widefield imaging causes out-of-focus light to appear in images of 3D samples. 2D SR-SIM increases the lateral resolution by extending the OTF along the *k*_*x*_ and *k*_*y*_ directions: the illumination pattern shifts information into the passband of the OTF in the direction determined by its orientation. By using three orientations, complete isotropic extension of the OTF can be achieved. In this case, the spatial frequency should be near the edge of the OTF in order to maximise the resolution increase. In the case of OS-SIM, the aim is to fill the missing cone. For this, the OTF must be shifted laterally in *k*_*z*_ so that the side lobes are centered on the origin, to cover the missing cone area in frequency space. For this, an illumination spatial frequency of approximately half the diffraction limit is required.

**Fig. 9.**
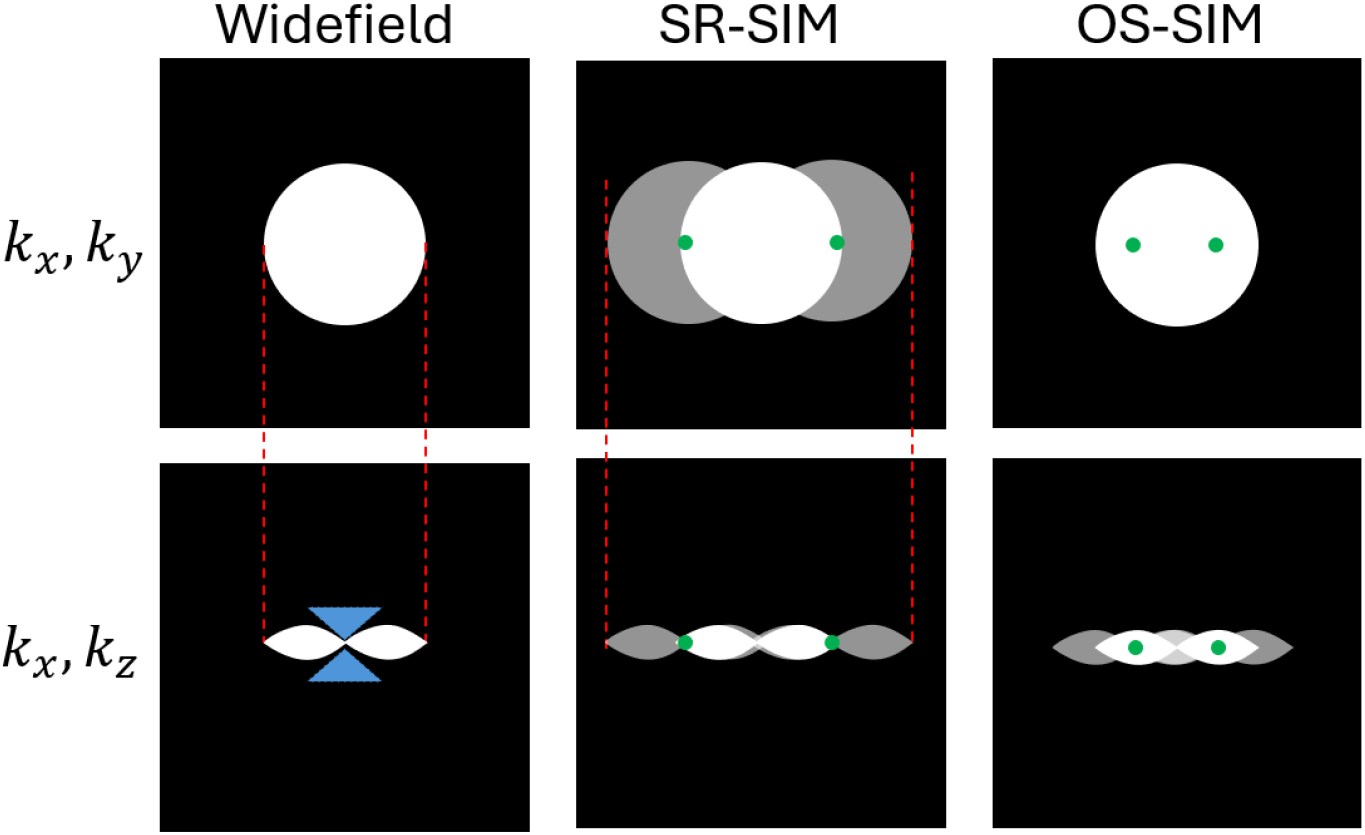
Illustration of the different spatial frequencies required for OS- and SR-SIM. The widefield OTF (white) can be extended to cover a greater area in frequency space (grey) by using striped illumination patterns, represented by a single *k*_*x,y*_ vector in frequency space (green). The two techniques improve the OTF in different ways: SR-SIM extends the OTF in *k*_*x*_ and *k*_*y*_ to improve the x and y resolution, OS-SIM fills the “missing cone” (blue) of unknown axial information.

## Appendix B Further optical setup details

### B.1 Fibre array

For the fibre array, a 9-channel array was chosen to provide a sufficient range of stripe pattern spacings in conjunction with a 30-mm collimating lens and various objectives. The fibre type is S405-XP, enabling use of the array across the 400 - 680 nm wavelength band. The spacing between the fibres was chosen to be 127 µm, the minimum spacing possible, in order to provide the finest control of the increments of stripe pattern spacings (see B.2). For a given focal length of tube lens, the number and spacing of the channels in the fibre array, as well as the focal length and diameter of the collimating lens, should be chosen together to enable the correct magnification of the beam spots from the fibre tips to the back focal plane of the objective, with sufficient adjustment over the full range of the field of view. In this work, a compact system, and hence a short optical path length between the array and the tube lens, was the priority, so a 1-inch diameter collimating lens with the shortest focal length was chosen, and the array matched accordingly.

### B.2 Switching between SIM modes

In order to switch between the different modes of SIM imaging, such as OS and SR (Fig. 9), the spatial frequency of the illumination patterns must be altered. This is achieved by controlling the position where the beams are incident on the BFP of the objective lens. Usually when outputting light from fibres, the separation of the fibre tips is fixed; the separation of the beams at the BFP is controlled by means of changing the configuration of lenses and mirrors in the path between the fibre outputs and the objective. This inflexible configuration requires moving and/or replacing components, then re-aligning them, every time the user changes objective lens or different imaging modes are needed. Use of a multichannel fibre array allows for quick adjustment by simply unplugging the fibres from the fibre circuit and reconnecting them to a different pair of inputs in the fibre array (Fig. 10). The array and all lenses remain fixed in place.

**Fig. 10.**
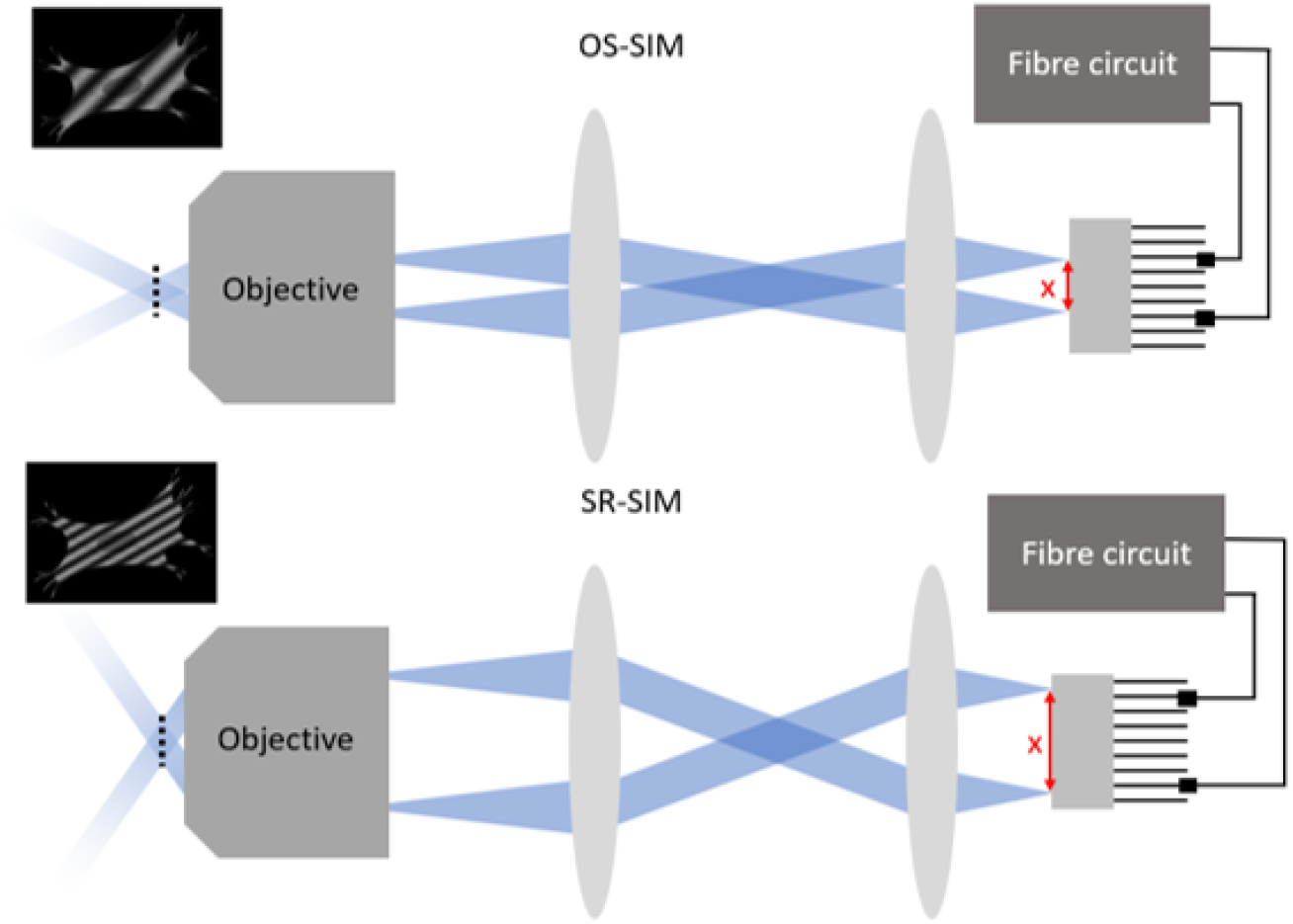
The use of a multichannel fibre array enables easy switching between different SIM modes and objective lenses, without changing, moving or re-aligning any of the optical components. Connections to different input fibres control the separation, x, of the spots at the face of the array, which in turn controls the separation at the BFP of the objective and hence the spatial frequency of the illumination patterns.

## Appendix C Resolution estimation

### C.1 Bead analysis

The resolution improvement was quantified for two illumination wavelengths by comparing the FWHM of sub-diffraction beads in diffraction-limited widefield and reconstructed SIM images.

The bead locations were extracted from images using peak local maxima, excluding beads that were within 2 µm of neighbouring beads. The FWHM was extracted by fitting a Gaussian function to the intensity profile of beads, and the resolution was subsequently estimated from the mean of the FWHM of beads after outliers were removed using a histogram of bead FWHMs.

### C.2 Decorrelation analysis

To quantify the resolution improvement of SR-SIM images, decorrelation analysis [32] was applied to widefield and reconstructed SIM images of immunostained *β−* tubulin in fixed Vero cells. Image analysis was performed using the Image Decorrelation plugin for ImageJ [32]. Results are shown in Fig. 11.

**Fig. 11.**
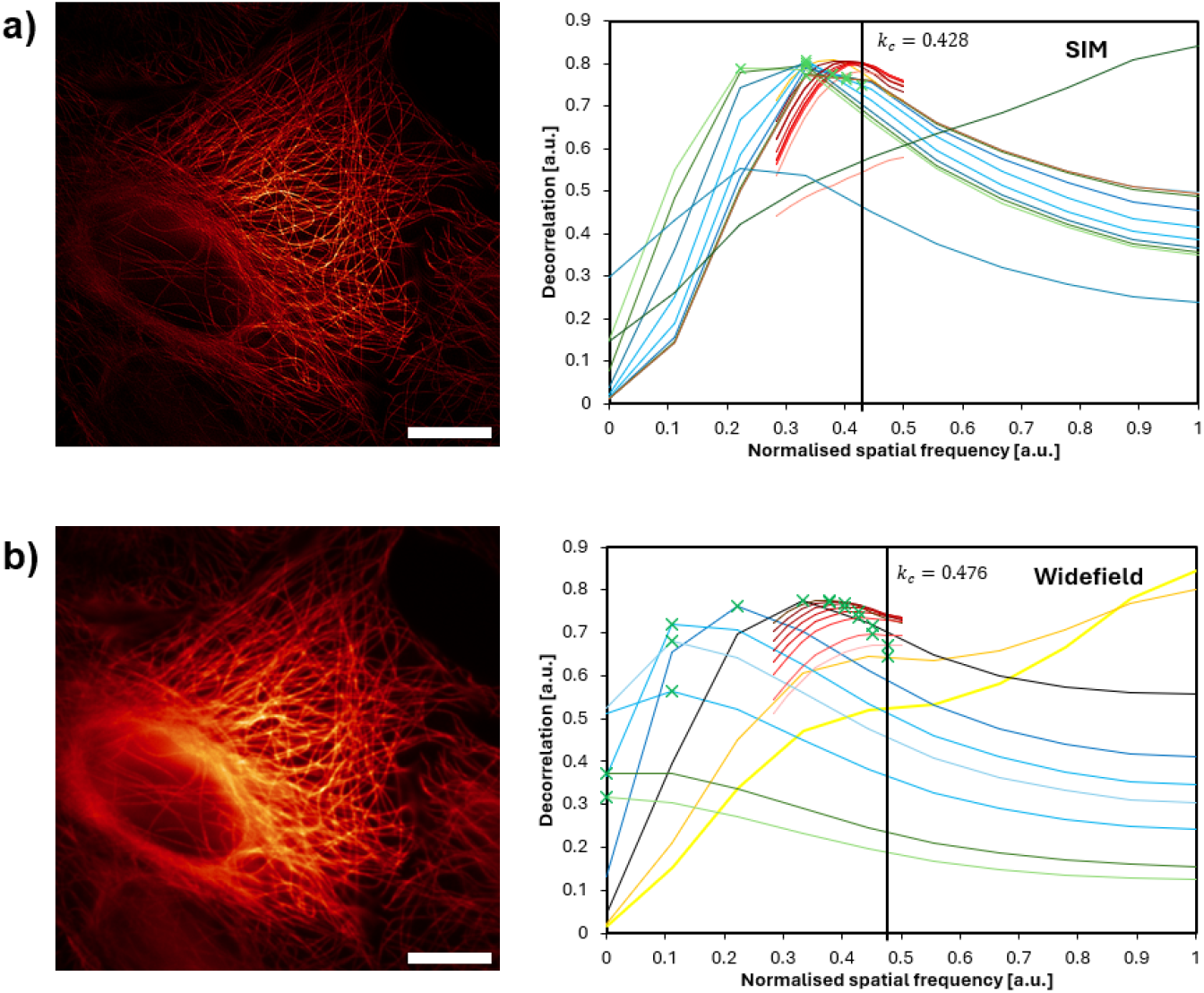
Decorrelation analysis estimates a resolution improvement from 306 to 201 nm when comparing a) SR-SIM and b) widefield images. Original images are shown on the left and their corresponding decorrelation curves on the right. (n=1) Scale bars are 10 µm. *k*_*c*_ is the cut-off frequency.

## Appendix D Sample preparation

### D.1 Fixed Vero cell culture

Vero cells (from monkey kidney tissue) were plated into 8-well plates (Ibidi), 20,000 cells per well, and cultured under standard conditions (37 ^°^C, 5% CO_2_) in minimum essential medium (Sigma Aldrich) supplemented with 10% foetal bovine serum (Gibco) and 2 mM L-Glutamine (GlutaMAX, Gibco). After 24 h, cells were fixed by incubation with 4% methanol-free formaldehyde and 0.1% glutaraldehyde in cacodylate buffer (pH 7.4) for 15 min at room temperature, washed three times with PBS and then permeabilised by incubation with a 0.2% solution of saponin in PBS for 15 min. Unspecific binding was blocked by incubating with 10% goat serum and 100 mM glycine in PBS and 0.2% saponin for 30 min at room temperature. Without washing, the samples were incubated with the primary antibody (mouse anti-beta-tubulin: ab131205) diluted 1:200 in PBS containing 2% BSA (bovine serum albumin) and 0.2% saponin overnight at 4 ^°^C. After three washes in PBS, the samples were incubated with the secondary antibody (goat anti-mouse conjugated to AlexaFluor568) diluted 1:400 in PBS containing 2% BSA and 0.2% saponin for 1 h at room temperature in the dark. Samples were then washed 3 times with PBS and imaged.

### D.2 Live COS-7 cell culture

**Cell culture** COS-7 cells were grown in T75 or T25 flasks by incubation at 37^°^C in a 5% CO_2_ atmosphere. Complete medium for normal cell growth consisted of 90% Dulbecco’s modified Eagle’s medium (DMEM) and 10% fetal bovine serum (FBS), supplemented with 1% GlutaMAX and 1% penicillin-streptomycin. Splitting was performed at >80% confluency and medium was refreshed every 3–4 days. For imaging, cells were seeded into 8-well plates (Ibidi) with 200 µL of standard culture medium.

**Labelling of organelles for live imaging** For ER labelling, COS-7 cells were transfected with plasmid constructs with Lipofectamine 2000 according to the manufacturer’s protocol 24 h before imaging. The plasmid used was pEGFPC1-hVAP-A (Addgene plasmid #104447), which was a gift from C. Tomasetto. For mitochondrion labelling, cells were incubated with MitoTracker Orange (Invitrogen, M7510) at 1 µM in culture medium for 15 min. Cells were then washed five times with culture medium and imaged immediately. Cells were imaged in a microscope stage-top incubator (OKOLab) with continuous air supply (37^°^C and 5% CO_2_).

### D.3 Bead monolayer

Monolayers of 0.1 µm fluorescent microspheres (ThermoFisher, T7279) and 0.3 µm microspheres labelled with Rhodamine B (courtesy of Andrew Harrison) were prepared in 8-well chambered coverglasses (Thermofisher, 155411), for resolution estimation and quantification of pattern spacing, respectively. Beads were first suspended in deionised water to a concentration of 2.310^5^ beads/ml. Bead monolayers were then formed by air drying 4 µl of sonicated bead dilutions in wells. Prior to imaging, 0.2 ml of deionised water was added to the wells to match the refractive index of the objective immersion media.

## References

[1] E. N. Ward, R. M. McClelland, J. R. Lamb, et al., “Fast, multicolour optical sectioning over extended fields of view with patterned illumination and machine learning,” Biomed. Opt. Express 15, 1074 (2024).

[2] M. A. A. Neil, R. Juškaitis, and T. Wilson, “Method of obtaining optical sectioning by using structured light in a conventional microscope,” Opt. Lett. 22, 1905–1907 (1997).

[3] R. Heintzmann and C. G. Cremer, “Laterally modulated excitation microscopy: improvement of resolution by using a diffraction grating,” in Optical Biopsies and Microscopic Techniques III, vol. 3568 (SPIE, 1999), pp. 185–196.

[4] M. G. L. Gustafsson, “Surpassing the lateral resolution limit by a factor of two using structured illumination microscopy,” J. Microsc. 198, 82–87 (2000).

[5] J. T. Frohn, H. F. Knapp, and A. Stemmer, “True optical resolution beyond the Rayleigh limit achieved by standing wave illumination,” Proc. National Acad. Sci. United States Am. 97, 7232–7236 (2000).

[6] G. E. Cragg and P. T. C. So, “Lateral resolution enhancement with standing evanescent waves,” Opt. Lett. 25 (2000).

[7] M. G. Gustafsson, “Nonlinear structured-illumination microscopy: Wide-field fluorescence imaging with theoretically unlimited resolution,” Proc. National Acad. Sci. United States Am. 102, 13081–13086 (2005).

[8] R. Heintzmann, “Saturated patterned excitation microscopy with two-dimensional excitation patterns,” in Micron, vol. 34 (Elsevier Ltd, 2003), pp. 283–291.

[9] D. Li, L. Shao, B. C. Chen, et al., “Extended-resolution structured illumination imaging of endocytic and cytoskeletal dynamics,” Science 349 (2015).

[10] M. Schropp and R. Uhl, “Two-dimensional structured illumination microscopy,” J. Microsc. 256, 23–36 (2014).

[11] E. Mudry, K. Belkebir, J. Girard, et al., “Structured illumination microscopy using unknown speckle patterns,” Nat. Photonics 6, 312–315 (2012).

[12] M. G. Gustafsson, L. Shao, P. M. Carlton, et al., “Three-dimensional resolution doubling in wide-field fluorescence microscopy by structured illumination,” Biophys. J. 94, 4957–4970 (2008).

[13] A. G. York, S. H. Parekh, D. D. Nogare, et al., “Resolution doubling in live, multicellular organisms via multifocal structured illumination microscopy,” Nat. Methods 9, 749–754 (2012).

[14] X. Huang, J. Fan, L. Li, et al., “Fast, long-term, super-resolution imaging with Hessian structured illumination microscopy,” Nat. Biotechnol. 36, 451–459 (2018).

[15] F. Ströhl and C. F. Kaminski, “Speed limits of structured illumination microscopy,” Opt. Lett. 42, 2511 (2017).

[16] E. N. Ward, L. Hecker, C. N. Christensen, et al., “Machine learning assisted interferometric structured illumination microscopy for dynamic biological imaging,” Nat. Commun. 13 (2022).

[17] E. N. Ward, C. N. Christensen, and R. M. McClelland, “ML-OS-SIM,” GitHub, https://github.com/edward-n-ward/ML-OS-SIM (2023).

[18] H. Zhang, Y. Zhu, L. Jin, et al., “Recent Advances in Structured Illumination Microscopy: From Fundamental Principles to AI-Enhanced Imaging,” (2025).

[19] X. Chen, S. Zhong, Y. Hou, et al., “Superresolution structured illumination microscopy reconstruction algorithms: a review,” Light. Sci. & Appl. 12, 172 (2023).

[20] L. J. Young, F. Ströhl, and C. F. Kaminski, “A guide to structured illumination TIRF microscopy at high speed with multiple colors,” J. Vis. Exp. 2016 (2016).

[21] A. Sandmeyer, M. Lachetta, H. Sandmeyer, et al., “Cost-Effective Live Cell Structured Illumination Microscopy with Video-Rate Imaging,” ACS Photonics 8, 1639–1648 (2021).

[22] D. Dan, M. Lei, B. Yao, et al., “DMD-based LED-illumination Super-resolution and optical sectioning microscopy,” Sci. Reports 3, 1116 (2013).

[23] P. Tinning, M. Donnachie, J. Christopher, et al., “Miniaturized structured illumination microscopy using two 3-axis MEMS micromirrors,” Biomed. Opt. Express 13, 6443 (2022).

[24] W. Liu, Q. Liu, Z. Zhang, et al., “Three-dimensional super-resolution imaging of live whole cells using galvanometer-based structured illumination microscopy,” Opt. Express 27, 7237 (2019).

[25] M. Brunstein, K. Wicker, K. Hérault, et al., “Full-field dual-color 100-nm super-resolution imaging reveals organization and dynamics of mitochondrial and ER networks,” Opt. Express 21, 26162 (2013).

[26] H. Ortkrass, J. Schürstedt, G. Wiebusch, et al., “High-speed TIRF and 2D super-resolution structured illumination microscopy with a large field of view based on fiber optic components,” Opt. Express 31, 29156 (2023).

[27] T. A. Hinsdale, S. Stallinga, and B. Rieger, “High-speed multicolor structured illumination microscopy using a hexagonal single mode fiber array,” Biomed. Opt. Express 12, 1181 (2021).

[28] S. Hu, W. Liu, J. Jie, et al., “Structured illumination microscopy based on fiber devices,” Opt. Commun. 460 (2020).

[29] J. Pospíšil, G. Wiebusch, K. Fliegel, et al., “Highly compact and cost-effective 2-beam super-resolution structured illumination microscope based on all-fiber optic components,” Opt. Express 29, 11833 (2021).

[30] M. Müller, V. Mönkemöller, S. Hennig, et al., “Open-source image reconstruction of super-resolution structured illumination microscopy data in ImageJ,” Nat. Commun. 7 (2016).

[31] R. Cao, Y. Chen, W. Liu, et al., “Inverse matrix based phase estimation algorithm for structured illumination microscopy,” Biomed. Opt. Express 9, 5037 (2018).

[32] A. Descloux, K. S. Grußmayer, and A. Radenovic, “Parameter-free image resolution estimation based on decorrelation analysis,” Nat. Methods 16, 918–924 (2019).

[33] G. Voeltz, E. Sawyer, G. Hajnóczky, and W. Prinz, “Making the connection: How membrane contact sites have changed our view of organelle biology,” Cell 187, 257–270 (2024).

[34] C. Stringer, T. Wang, M. Michaelos, and M. Pachitariu, “Cellpose: a generalist algorithm for cellular segmentation,” Nat. Methods 18, 100–106 (2021).

[35] A. E. Carpenter, T. R. Jones, M. R. Lamprecht, et al., “CellProfiler: image analysis software for identifying and quantifying cell phenotypes,” Genome Biol. 7, R100 (2006).

[36] A. E. Y. T. Lefebvre, G. Sturm, T.-Y. Lin, et al., “Nellie: automated organelle segmentation, tracking and hierarchical feature extraction in 2D/3D live-cell microscopy,” Nat. Methods 22, 751–763 (2025).

[37] S. Munck, C. De Bo, C. Cawthorne, and J. Colombelli, “Innovating in a bioimaging core through instrument development,” J. Microsc. 294, 319–337 (2024).

[38] M. Weber and J. Huisken, “Multidisciplinarity Is Critical to Unlock the Full Potential of Modern Light Microscopy,” Front. Cell Dev. Biol. 9 (2021).

[39] H. Wang, P. T. Brown, J. Ullom, et al., “Fully automated multicolour structured illumination module for super-resolution microscopy with two excitation colours,” Commun. Eng. 4 (2025).

[40] M. T. Hannebelle, E. Raeth, S. M. Leitao, et al., “Open-source microscope add-on for structured illumination microscopy,” Nat. Commun. 15 (2024).

[41] H. Wang, R. Lachmann, B. Marsikova, et al., “UCsim2: two-dimensionally structured illumination microscopy using UC2,” Philos. Trans. Royal Soc. A: Math. Phys. Eng. Sci. 380 (2022).

[42] R. M. Power and J. Huisken, “Putting advanced microscopy in the hands of biologists,” Nat. Methods 16, 1069–1073 (2019).

[43] R. M. Power, T. Bakken, J. Li, and J. Huisken, “Expanding advanced microscopy access with a customizable and shareable light sheet microscope (Conference Presentation),” in High-Speed Biomedical Imaging and Spectroscopy IV, K. Goda and K. K. Tsia, eds. (SPIE, 2019), p. 23.

[44] D. L. Coutu and T. Schroeder, “Probing cellular processes by long-term live imaging – historic problems and current solutions,” J. Cell Sci. 126 (2013).

[45] H. Yan, T. Wu, X. Li, et al., “Establishment of the microscope incubation system and its application in evaluating tumor treatment effects through real-time live cellular imaging,” Front. Bioeng. Biotechnol. 12 (2024).

[46] B. Mandracchia, C. Zheng, S. Rajendran, et al., “High-speed optical imaging with sCMOS pixel reassignment,” Nat. Commun. 15 (2024).

[47] A. A. Pulschen, D. R. Mutavchiev, S. Culley, et al., “Live Imaging of a Hyperthermophilic Archaeon Reveals Distinct Roles for Two ESCRT-III Homologs in Ensuring a Robust and Symmetric Division,” Curr. Biol. 30, 2852–2859 (2020).

[48] M. Nishiyama, “High-pressure microscopy for tracking dynamic properties of molecular machines,” Biophys. Chem. 231, 71–78 (2017).

[49] S. Hupp, N. S. Tomov, C. Bischoff, et al., “Easy to build cost-effective acute brain slice incubation system for parallel analysis of multiple treatment conditions,” J. Neurosci. Methods 366, 109405 (2022).

[50] A. Talebipour, M. Saviz, M. Vafaiee, and R. Faraji-Dana, “Facilitating long-term cell examinations and time-lapse recordings in cell biology research with CO2 mini-incubators,” Sci. Reports 14, 3418 (2024).

[51] A. P. M. Marty, E. N. Ward, J. R. Lamb, et al., “A High-Resolution Microscopy System for Biological Studies of Cold-Adapted Species Under Physiological Conditions,” Small Methods 9 (2025).

[52] C. W. Chung, A. D. Stephens, T. Konno, et al., “Intracellular Aβ42 Aggregation Leads to Cellular Thermogenesis,” J. Am. Chem. Soc. 144, 10034–10041 (2022).

[53] G. O. Merces, C. Kennedy, B. Lenoci, et al., “The incubot: A 3D printer-based microscope for long-term live cell imaging within a tissue culture incubator,” HardwareX 9, e00189 (2021).

[54] D. Jin, D. Wong, J. Li, et al., “Compact Wireless Microscope for In-Situ Time Course Study of Large Scale Cell Dynamics within an Incubator,” Sci. Reports 5 (2015).

[55] S. B. Kim, K.-i. Koo, H. Bae, et al., “A mini-microscope for in situ monitoring of cells.” Lab on a chip 12, 3976–3982 (2012).

[56] M. P. Walzik, V. Vollmar, T. Lachnit, et al., “A portable low-cost long-term live-cell imaging platform for biomedical research and education,” Biosens. Bioelectron. 64, 639–649 (2015).

