## Supplementary figures and images for "SIMple: A fibre-based platform for accessible structured illumination microscopy"

### Supplemental Video 1

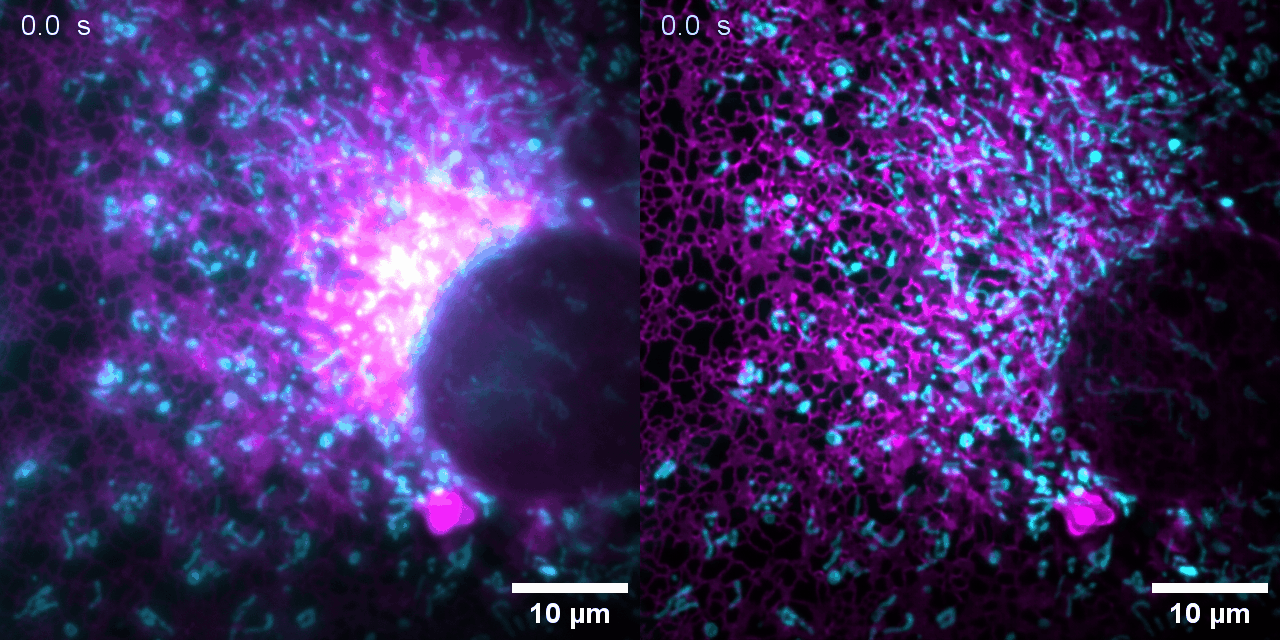
